# Adequate knowledge of COVID-19 impacts good practices amongst health profession students in the Philippines

**DOI:** 10.1101/2021.02.18.431919

**Authors:** Junhel Dalanon, Rhomeljustein Redoble, Jo-Ann Belotindos, Candy Delos Reyes, Jaime Fabillar, Ma. Shiril Armero, Rozzano Locsin, Yoshitaka Suzuki, Kazuo Okura, Yoshizo Matsuka

**Author notes:** Corresponding author at: School of Dentistry, Southwestern University PHINMA, Cebu, Philippines 6000.

## Abstract

**Background:** The spread of the coronavirus disease 2019 (COVID-19) in the Philippines started with its first suspected case on January 22, 2020. The government reacted by imposing several measures including community quarantine, class suspensions, drug therapy and vaccine development, and travel restrictions. This online survey was done amongst Filipino health professions undergraduate students to uncover the relationship between their knowledge, attitude, and practice during this pandemic.

**Methods:** Cross-sectional data were obtained from an online survey done on students of medicine, dentistry, optometry, rehabilitative sciences, and pharmacy.

**Results:** At a response rate of 100% (n=1257), the results show that healthcare profession students in the Philippines have good knowledge (87.6%) and practices (63.6%) regarding COVID-19, yet attitude (63.6%) was just passable. This study also shows that a strong correlation exists between knowledge and practice concerning the current pandemic, r(2) = 0.08, P = 0.004.

**Conclusion:** Adequate knowledge of COVID-19 impacts good practices of avoiding crowded places and misuse of steam inhalation amongst health profession students in the Philippines. Knowledge and practice pertaining to the current pandemic have been found to be good, but attitude remains low.

## 1. Introduction

Since the end of 2019, the attention of the scientific community has been on the COVID-19 pandemic which originated from Wuhan City, China and which is caused by the SARS-CoV-2 [1]. Belonging to the Coronaviridae, the coronaviruses contain a single strand of RNA (26-32 kb) and a diameter of 65 to 125 nm. Alpha, beta, gamma, delta, AH1N1, MERS-CoV, H5N1, influenza A, and SARS-CoV comprises the coronavirus types [2, 3]. On February 11, 2020, the International Committee on Taxonomy of Viruses (ICTV) named the disease as COVID-19 and the virus as SARS-CoV-2 [4, 5].

There have been 95, 544, 853 cases registered worldwide and 2,039, 947 deaths globally [6]. In the Philippines, there have been 500, 577 confirmed cases, 9, 985 deaths, and 1, 886 new cases according to the World Health Organization (WHO) [7]. Deemed to be equally a life-threatening disease and an exceedingly contagious disease, the WHO has classified COVID-19 as a pandemic. The expected number of cases generated by a single infected individual within a susceptible population [8] or the reproduction ratio (RR) of the disease (RR = 2.4-2.7) are greater compared to influenza (RR = 1.3-1.8) [9, 10].

From a severe ailment with extreme death rate to an infection with no symptoms, the clinical manifestation is wide-ranging. Dry coughing, fever, and fatigue that can predispose to chest pains, difficulty in breathing, moving, and talking are some of the common symptoms of the disease [4, 11]. Transmission of SARS-CoV-2 may be through indirect contact with contaminated items or human-to-human contact [12, 13]. In humans with COVID-19, mouth or nose transmission of body fluid droplets have been documented during talking, coughing, or sneezing. Despite being able to traverse only within six feet, the virus can be suspended in the air for up to three hours and can remain intact in droplets [14]. SARS-CoV-2 can be transmitted through contaminated cardboard, stainless steel, copper, or plastic to the mucous membranes of the nose, mouth, or eyes [15].

It has been established that the knowledge, attitude, and practice (KAP) of a specific population are necessary indices to avoid distress and panic. Surveys on KAP can likewise provide insights on what is being believed, done, and known by the community. This becomes very important in these times of crisis [16–18].

## 2. Subjects and methods

### 2.1. Study design and population

This is an intercollegiate cross-sectional study conducted in the School of Medicine, School of Dentistry, School of Optometry, College of Pharmacy, and College of Rehabilitative Sciences in a higher education institution in the Philippines from July 7, 2020 to December 31, 2020. The participants were purposively sampled students from these aforementioned colleges.

### 2.2. Study instrument

After comprehensive literature search, a modified instrument was devised from a pre-validated questionnaire [17]. Information from the Department of Health of the Philippines, the WHO, and the Center for Disease Control and Prevention were also considered. As the Philippines uses as the medium of instruction in its basic education and higher education schools, the authors did not deem it necessary to translate and adapt the language. Data gathering was done using a readily accessible online survey software. The questionnaire contained 4 sections. The first section was composed of the demographic profile. The second section contained 12 questions that gauged the knowledge of the participants about COVID-19. The third section evaluated their attitude on (A1) optimism in successfully controlling the pandemic, (A2) confidence that the Philippines will be able to control the virus, (A3) hope that their family can withstand this ordeal, and (A4) positivity that they can still enroll for the next school year. The last section had 4 questions that covered COVID-19 related practices. These questions weighed-in on how they (P1) avoided crowded places, (P2) if they wore masks, (P3) washed their hands, and (A4) avoided the misuse of steam inhalation as a cure for the disease.

### 2.3. Ethical considerations

Ethical approval was sought from and given by the ethical review board of the School of Dentistry of Southwestern University PHINMA. Prior to the start of data gathering, the participants were oriented with the goal of the study, informed that their identities will be maintained anonymous, that their responses will not affect their academic performance in any way, and that they could opt out of the study at any time. Digital signatures of the participants were collected as consent to participate in the study.

### 2.4. Statistical analysis

Data obtained were coded and analyzed using the Statistical Package for Social Science, version 26.0 (IBM Corp., Armonk, NY) Frequencies and proportions were calculated using descriptive analysis by cross-tabulation and using chi-square test for association. Spearman’s correlation was used for nonparametric measure of rank correlation. Logistic regression analysis was used to detect risk factors linked to attitudes and practices. The normality of data was tested using the Kolmogorov-Smirnov test. In this study, a p-value of less than 0.05 was considered statistically significant.

## 3. Results

With a 100% completion rate, a total of 1257 health profession students were able to finish all the sections of the survey questionnaire. There were 1031 (82%) medical students, 76 (6%) students of dentistry, 80 (6.4%) optometry students, 18 (1.4%) students from the rehabilitative sciences, and 51 (4.1%) pharmacy students.

In terms of knowledge, significant associations were found with age (*X*^2^_(3, N = 1257)_ = 8.66, P = 0.034), gender (*X*^2^_(9, N = 1257)_ = 24.64, P = 0.003), citizenship (*X*^2^_(1, N = 1257)_ = 16.6, P < 0.001), year level (*X*^2^_(6, N = 1257)_ = 34.17, P < 0.001), place of residence (*X*^2^_(3, N = 1257)_ = 13.51, P = 0.004), and quarantine conditions (*X*^2^_(3, N = 1257)_ = 14.38, P = 0.002). Those who showed more knowledge in the context of COVID-19 were 25-30 years old (91.9%), females (90%), Filipinos (90.3%), past their 6^th^ year in college (100%), living near the university (90.4%), and subjected to MECQ conditions (92.1%). Moreover, attitude was suggestively associated with age (*X*^2^_(3, N = 1257)_ = 15.84, P = 0.001), gender (*X*^2^_(9, N = 1257)_ = 30.74, P < 0.001), citizenship (*X*^2^_(1, N = 1257)_ = 129.74, P < 0.001), college program (*X*^2^_(4, N = 1257)_ = 42.59, P < 0.001), and year level (*X*^2^_(6, N = 1257)_ = 20.12, P = 0.003). In which health profession students more than 30 years old (83%), males (71.3%), foreigners (85.3%), belonging to the college of medicine (67.7%), and past their 6^th^ year of college (100%) were more optimistic about the current pandemic. On the other hand, good practices that help thwart the threat of COVID-19 were considerably linked with year level (*X*^2^_(6, N = 1257)_ = 14.65, P = 0.023). The data suggests that students in their 5^th^ year (100%) and those who spent beyond 6 years of college (100%) yielded better practices against the current pandemic (Table 1). Using Spearman’s rank correlation, a strong correlation was found between knowledge and practice, *r*(2) = 0.08, P = 0.004 (Table 2).

**Table 1.**
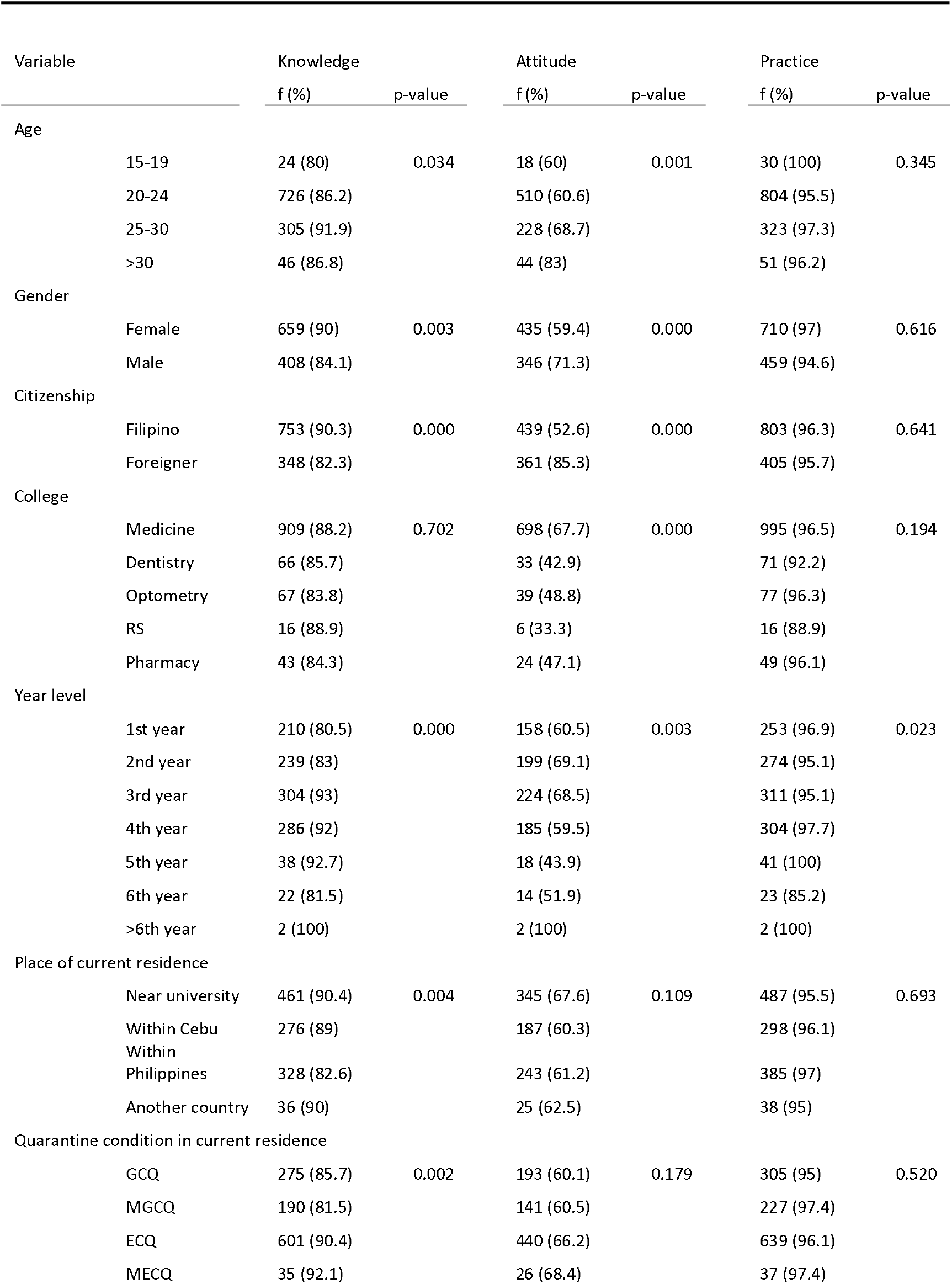
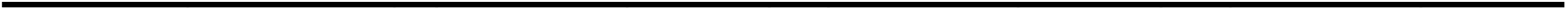
Cross-tabulation of the sociodemographic characteristics of health professions students with knowledge, attitude, and practice

| Variable |  | Knowledge |  | Attitude |  | Practice |  |
| --- | --- | --- | --- | --- | --- | --- | --- |
|  |  | f (%) | p-value | f (%) | p-value | f (%) | p-value |
| Age |  |  |  |  |  |  |  |
|  | 15-19 | 24 (80) | 0.034 | 18 (60) | 0.001 | 30 (100) | 0.345 |
|  | 20-24 | 726 (86.2) |  | 510 (60.6) |  | 804 (95.5) |  |
|  | 25-30 | 305 (91.9) |  | 228 (68.7) |  | 323 (97.3) |  |
|  | >30 | 46 (86.8) |  | 44 (83) |  | 51 (96.2) |  |
| Gender |  |  |  |  |  |  |  |
|  | Female | 659 (90) | 0.003 | 435 (59.4) | 0.000 | 710 (97) | 0.616 |
|  | Male | 408 (84.1) |  | 346 (71.3) |  | 459 (94.6) |  |
| Citizenship |  |  |  |  |  |  |  |
|  | Filipino | 753 (90.3) | 0.000 | 439 (52.6) | 0.000 | 803 (96.3) | 0.641 |
|  | Foreigner | 348 (82.3) |  | 361 (85.3) |  | 405 (95.7) |  |
| College |  |  |  |  |  |  |  |
|  | Medicine | 909 (88.2) | 0.702 | 698 (67.7) | 0.000 | 995 (96.5) | 0.194 |
|  | Dentistry | 66 (85.7) |  | 33 (42.9) |  | 71 (92.2) |  |
|  | Optometry | 67 (83.8) |  | 39 (48.8) |  | 77 (96.3) |  |
|  | RS | 16 (88.9) |  | 6 (33.3) |  | 16 (88.9) |  |
|  | Pharmacy | 43 (84.3) |  | 24 (47.1) |  | 49 (96.1) |  |
| Year level |  |  |  |  |  |  |  |
|  | 1st year | 210 (80.5) | 0.000 | 158 (60.5) | 0.003 | 253 (96.9) | 0.023 |
|  | 2nd year | 239 (83) |  | 199 (69.1) |  | 274 (95.1) |  |
|  | 3rd year | 304 (93) |  | 224 (68.5) |  | 311 (95.1) |  |
|  | 4th year | 286 (92) |  | 185 (59.5) |  | 304 (97.7) |  |
|  | 5th year | 38 (92.7) |  | 18 (43.9) |  | 41 (100) |  |
|  | 6th year | 22 (81.5) |  | 14 (51.9) |  | 23 (85.2) |  |
|  | >6th year | 2 (100) |  | 2 (100) |  | 2 (100) |  |
| Place of current residence |  |  |  |  |  |  |  |
|  | Near university | 461 (90.4) | 0.004 | 345 (67.6) | 0.109 | 487 (95.5) | 0.693 |
|  | Within Cebu | 276 (89) |  | 187 (60.3) |  | 298 (96.1) |  |
|  | Within Philippines | 328 (82.6) |  | 243 (61.2) |  | 385 (97) |  |
|  | Another country | 36 (90) |  | 25 (62.5) |  | 38 (95) |  |
| Quarantine condition in current residence |  |  |  |  |  |  |  |
|  | GCQ | 275 (85.7) | 0.002 | 193 (60.1) | 0.179 | 305 (95) | 0.520 |
|  | MGCQ | 190 (81.5) |  | 141 (60.5) |  | 227 (97.4) |  |
|  | ECQ | 601 (90.4) |  | 440 (66.2) |  | 639 (96.1) |  |
|  | MECQ | 35 (92.1) |  | 26 (68.4) |  | 37 (97.4) |  |

**Table 2.** Correlation between knowledge, attitude, and practice

| Variables | rho | p-value |
| --- | --- | --- |
| Knowledge and attitude | 0.007 | 0.799 |
| Knowledge and practice | 0.081 | 0.004** |
| Attitude and practice | 0.048 | 0.09 |

In terms of attitude, results of the regression analysis found that being Filipino is a factor in being optimistic that the Philippines will win the battle against the pandemic (OR: 0.248, P<0.0001), that their respective families will overcome the hardships (OR: 0.203, P<0.001), and that they can still enroll next school year (OR: 0.449, P<0.0001). Moreover, being a medical student is a factor in believing that the pandemic will be successfully controlled (OR: 2.631, P<0.0001), and that enrollment in their programs next year is still an option (OR: 3.653, P<0.0001). Belonging to the 1^st^ year to 4^th^ year also is a factor in the optimism that the pandemic can be controlled (OR: 3.506, P<0.0001), and that they can continue to study next school year (OR: 1.719, P<0.05). In addition, living within Cebu City influences the health profession students’ confidence that the pandemic will be contained soon (OR: 1.398, P<0.05). Furthermore, those students quarantined other than in GCQ conditions were more enthusiastic that the pandemic will be resolved (OR: 0.591, P<0.001), their family will survive this ordeal (OR: 1.766, P<0.05), and that they will still be able to enroll next school year (OR: 0.654, P<0.001). Comparatively, the results show that in terms of practices being knowledgeable influences the practice of avoiding crowded places (OR: 2.978, P<0.0001) and avoiding the incorrect use of steam inhalation as a cure (OR: 1.112, P<0.05). Being a Filipino was found to be a factor that affects the health profession students to avoid crowded places (OR: 2.291, P<0.0001) and wash their hands (OR: 7.396, P<0.0001). Finally, living within Cebu City influences the avoidance of using steam inhalation as a cure for COVID-19 (OR: 1.878, P<0.001) (Table 3).

**Table 3.**
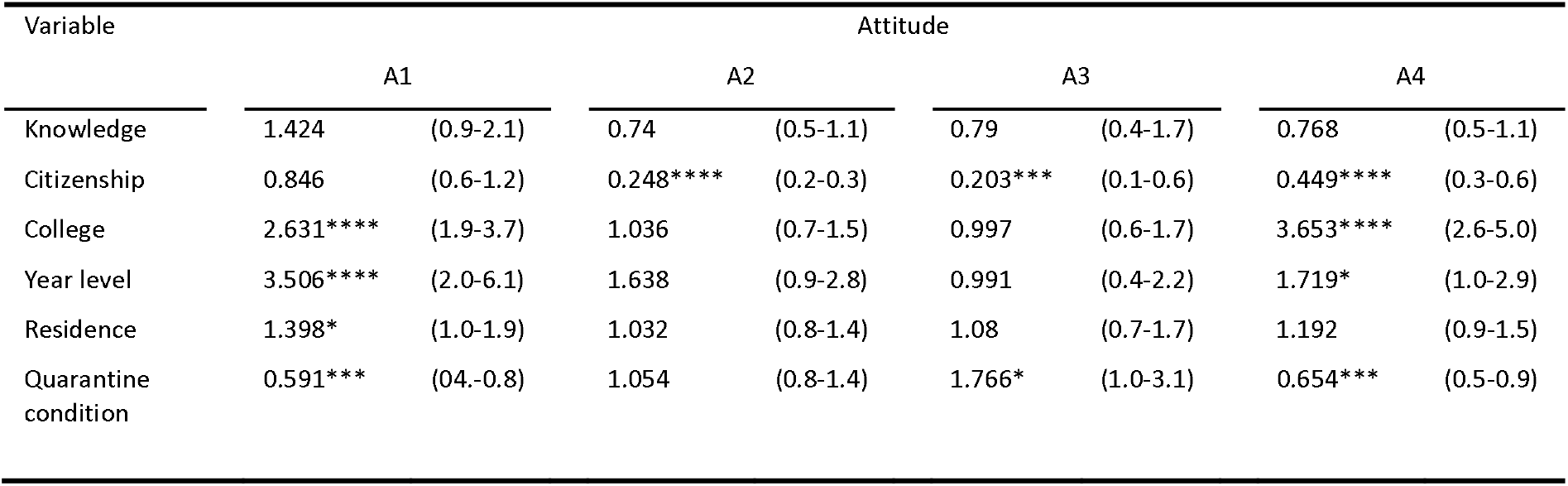
Significantly associated factors towards COVID-19 logistic regression analysis of odds ratio (OR) for attitudes in relation to potential risk factors

| Variable | Attitude |  |  |  |  |  |  |  |
| --- | --- | --- | --- | --- | --- | --- | --- | --- |
|  | A1 |  | A2 |  | A3 |  | A4 |  |
| Knowledge | 1.424 | (0.9-2.1) | 0.74 | (0.5-1.1) | 0.79 | (0.4-1.7) | 0.768 | (0.5-1.1) |
| Citizenship | 0.846 | (0.6-1.2) | 0.248**** | (0.2-0.3) | 0.203*** | (0.1-0.6) | 0.449**** | (0.3-0.6) |
| College | 2.631**** | (1.9-3.7) | 1.036 | (0.7-1.5) | 0.997 | (0.6-1.7) | 3.653**** | (2.6-5.0) |
| Year level | 3.506**** | (2.0-6.1) | 1.638 | (0.9-2.8) | 0.991 | (0.4-2.2) | 1.719* | (1.0-2.9) |
| Residence | 1.398* | (1.0-1.9) | 1.032 | (0.8-1.4) | 1.08 | (0.7-1.7) | 1.192 | (0.9-1.5) |
| Quarantine condition | 0.591*** | (0.4-0.8) | 1.054 | (0.8-1.4) | 1.766* | (1.0-3.1) | 0.654*** | (0.5-0.9) |

**Table 4.** Significantly associated factors towards COVID-19 logistic regression analysis of odds ratio (OR) for practices in relation to potential risk factors

| Variable | Practice |  |  |  |  |  |  |  |
| --- | --- | --- | --- | --- | --- | --- | --- | --- |
|  | P1 |  | P2 |  | P3 |  | P4 |  |
| Knowledge | 0.654 | (0.4-1.0) | 4.79 | (2.1-11.0) | 2.978**** | (1.0-8.7) | 1.112* | (0.7-1.8) |
| Citizenship | 2.291**** | (1.5-3.4) | 7.396**** | (2.8-19.9) | 0.611 | (0.2-2.3) | 0.939 | (0.6-1.4) |
| College | 0.686 | (0.5-1.0) | 0.536 | (0.2-1.7) | 1.525 | (0.5-4.7) | 0.908 | (0.6-1.4) |
| Year level | 1.146 | (0.6-2.3) | 1.288 | (0.3-5.9) | 0.71 | (0.1-5.8) | 0.672 | (0.4-1.3) |
| Residence | 0.918 | (0.7-1.3) | 1.6 | (0.7-3.6) | 0.862 | (0.3-2.4) | 1.878*** | (1.3-2.7) |
| Quarantine condition | 1.106 | (0.8-1.6) | 1.119 | (0.5-2.7) | 0.444 | (0.2-1.2) | 0.852 | (0.6-1.2) |

## 4. Discussion

On January 30, 2020, the first case of the coronavirus in the Philippines was reported when a 38-year-old Chinese woman from Wuhan-China was confirmed to be positive for the infectious disease [19]. Shortly thereafter, the country recorded its first local death on March 11, 2020 when a 67-year-old Filipina died from the disease [20]. On April 30, President Rodrigo Duterte placed the National Capital Region, Central Luzon, CALABARZON, Benguet, Pangasinan, Iloilo, Bacolod City, Davao City, and Cebu under Enhanced Community Quarantine (ECQ). The rest of the country was placed under General Community Quarantine (GCQ) [21]. With 4, 562 recorded cases of COVID-19 on June 29, 2020, Cebu City became the new epicenter in the country. In the ECQ no travel is allowed regardless of health status and age, economic activities are held to a minimum, private transportations are halted, and face-to-face classes are suspended. On the other hand, the GCQ allows limited travel to services and work, allows 75% workforce in government offices, permits limited transportation, and flexible classes are approved. Modifications of these quarantine classifications have also been put into place, where the modified ECQ (MECQ) and the modified GCQ (MGCQ) have been introduced. The MECQ is a more lenient version of the ECQ where limited travel for services and work are permitted, 50% workforce for manufacturing plants are allowed, limited transportation is approved, but physical classes are still suspended. Contrarily, socio-economic activities are tolerable with some public health standards. Provisions like these may limit the influx of information to the community [22].

To the knowledge of the authors, this is the first study to gauge the KAP of multiple categories of health profession students in a developing country towards COVID-19. Majority or 87.6% of the health profession students garnered good knowledge regarding COVID-19. This outcome is higher than the results from the knowledge of COVID-19 in Pakistani young adults (80%) [23] and Saudi University students on MERS-CoV (32%) [24]. However, this mark is slightly lower compared to the score from a study done in China (90%). The high results were attributed to the highest educational attainment of the respondents. They posited that the Chinese public has a wide array of information sources like the WeChat account of the Wuhan Health Commission, China Central Television, and the National Health Commission of China website [17]. This is similar to a study done on Italian undergraduate students of life science. They have attributed the high scores to sources of information like the television and the internet [25]. The data in this study suggests that older students, Females, Filipinos, who spent more than 6 years in college, living near the university, and under MECQ conditions are more knowledgeable about COVID-19. These findings are consistent with previous studies that found higher knowledge concerning COVID-19 as age progresses and in females [17, 25]. In the Philippine context, age and number of years in college might increase knowledge due to the structure of the educational system. In a Filipino family, females are also favored to go to universities as opposed to males who are expected to work [26]. Proximity to the university and being Filipino are also significant in terms in knowledge acquisition as teachers in healthcare professions can provide valuable information better [27]. In addition, language could also be a factor as some disease prevention information might be in the local language [28].

At 63.6%, the number of students who were optimistic of the current situation was just slightly more than half according to the results of the survey. This suboptimal optimism of the participants is opposite that of a couple of Chinese studies. They reasoned that the high confidence of Chinese residents was due to their government’s exceptional countermeasures, concerted efforts and ample resources, and good knowledge about COVID-19 [17]. At 87%, the attitude of Egyptian senior pharmacy students is also exceedingly higher than the results here [29]. The Philippines lack the experts, materials, and readiness in managing this epidemic. As a developing nation, the low economy could play a role in the effectiveness of managing a pandemic. This study found that older students, males, foreigners, enrolled in a medical program, and those who have spent more than 6 years in college are more optimistic in their attitude towards COVID-19. According to the results of the survey in terms of attitude, being Filipino influences optimism that the Philippines will win the war against the pandemic, that their families will survive this ordeal, and that they will still be able to pursue their education next school year. Being a medical student increases the confidence that the pandemic will be controlled soon and that they can still enroll for next school year. Students within their first four years of college are more hopeful that the coronavirus will be controlled, and they will be able to resume studying. Living in Cebu City is linked to the attitude of optimism that the virus will be contained soon. Students under GCQ are reported to be hopeful that the current pandemic will be resolved, that their families will survive unscathed, and they can continue enrolling in their programs. A study done among healthcare workers in Portugal found females with previous anxiety experience were less optimistic [30]. Being Filipino might also be key in the attitude regarding COVID-19 but care must be done in the interpretation of the results. A previous survey found that Filipinos are among the most confident, yet among the most ignorant in key issues. Nevertheless, previous studies have found that older, male, with right-wing political attitudes, living in urban areas have higher perceived resilience. This was credited to experiences of older individuals contributing to the development of greater confidence. In addition, Filipinos have a tendency to believe in a higher being and cling to religion in times of adversity [31]. The GCQ is the least restrictive of all of the quarantine measures category. Having preventive measures that are less hampering might have an effect on the optimism of the students. The perception of threat and the feeling of distress can lower the resilience of the community [32].

An overwhelming 96.1% of the participants perceived that they followed good practices in the prevention of COVID-19. This outcome is consistent with previous studies. Moreover, the findings convey that the more years spent in college, the more it translates in practicing precautionary measures against the disease. In terms of practices, having good knowledge impacts the habits of avoiding crowded places and the use of steam inhalation as a cure for the disease. Likewise, Filipinos are more inclined to avoid crowded places and to practice washing their hands. Living within the confines of Cebu City has an effect in evading the misuse of steam inhalation as a cure for COVID-19. The containment of a pandemic and rendering its consequences attenuated are dependent on the influence of KAP on the community’s adherence to preventive measures. The level of knowledge has also been linked to maintaining good practices towards the pandemic [33]. Perhaps a unique facet of this study is the discovery that living within Cebu City influences the avoidance of misusing steam inhalation as a cure for COVID-19. Steam inhalation has been controversial in this part of the Philippines when news broke that the city government spent 2.5 million pesos or 52000 US dollars’ worth of steam inhalation kits and was set to institutionalize steam inhalation. This was done despite several forewarnings from the local medical societies and the Department of Health [34].

The current study is not all-encompassing and there are several limitations that have been left unrequited. As with previous studies on KAP concerning COVID-19, majority of the participants were females. The socioeconomic status of the individuals would have been an excellent variable in determining how it affects KAP. Religion and how it affects the aforementioned variables can be an interesting perspective to unravel as well. As this study utilized online survey, there are still some sectors of the Philippine populace that don’t have online presence. New knowledge regarding the disease and its variants can modify the current findings. Multiple attitudes and practices can also be added in the future. More studies are warranted in investigating the KAP of healthcare professions students.

## 5. Conclusion

The findings of this study found high level of COVID-19 related knowledge among health profession students from a developing country. In addition, a positive association between knowledge and practices was found. Being older, a Filipino, a medical student, and in the first four years of college influences favorable attitude concerning COVID-19. Finally, good knowledge, being Filipino, and living within Cebu City improves the likelihood of adapting to good practices against the pandemic.

## References

[1] Fauci AS, Lane HC, Redfield RR. Covid-19 - Navigating the Uncharted. N Engl J Med. 2020;382(13):1268–9.

[2] Fehr AR, Perlman S. Coronaviruses: an overview of their replication and pathogenesis. Methods Mol Biol. 2015;1282:1–23.

[3] Pal M, Berhanu G, Desalegn C, Kandi V. Severe Acute Respiratory Syndrome Coronavirus-2 (SARS-CoV-2): An Update. Cureus. 2020;12(3):e7423.

[4] Huang C, Wang Y, Li X, Ren L, Zhao J, Hu Y, et al. Clinical features of patients infected with 2019 novel coronavirus in Wuhan, China. Lancet. 2020;395(10223):497–506.

[5] Wang D, Hu B, Hu C, Zhu F, Liu X, Zhang J, et al. Clinical Characteristics of 138 Hospitalized Patients With 2019 Novel Coronavirus-Infected Pneumonia in Wuhan, China. JAMA. 2020;323(11):1061–9.

[6] Dong E, Du H, Gardner L. An interactive web-based dashboard to track COVID-19 in real time. Lancet Infect Dis. 2020;20(5):533–4.

[7] World Health Organization. Philippines: WHO Coronavirus Disease (COVID-19) Dashboard Geneva: WHO; 2021 [updated January 19, 2021. Available from: https://covid19.who.int/region/wpro/country/ph.

[8] Fraser C, Donnelly CA, Cauchemez S, Hanage WP, Van Kerkhove MD, Hollingsworth TD, et al. Pandemic potential of a strain of influenza A (H1N1): early findings. Science. 2009;324(5934):1557–61.

[9] Zhao S, Lin Q, Ran J, Musa SS, Yang G, Wang W, et al. Preliminary estimation of the basic reproduction number of novel coronavirus (2019-nCoV) in China, from 2019 to 2020: A data-driven analysis in the early phase of the outbreak. Int J Infect Dis. 2020;92:214–7.

[10] Liu Y, Gayle AA, Wilder-Smith A, Rocklov J. The reproductive number of COVID-19 is higher compared to SARS coronavirus. J Travel Med. 2020;27(2).

[11] Xu XW, Wu XX, Jiang XG, Xu KJ, Ying LJ, Ma CL, et al. Clinical findings in a group of patients infected with the 2019 novel coronavirus (SARS-Cov-2) outside of Wuhan, China: retrospective case series. BMJ. 2020;368:m606.

[12] Chen Y, Liu Q, Guo D. Emerging coronaviruses: Genome structure, replication, and pathogenesis. J Med Virol. 2020;92(4):418–23.

[13] Lotfi M, Hamblin MR, Rezaei N. COVID-19: Transmission, prevention, and potential therapeutic opportunities. Clin Chim Acta. 2020;508:254–66.

[14] van Doremalen N, Bushmaker T, Morris DH, Holbrook MG, Gamble A, Williamson BN, et al. Aerosol and Surface Stability of SARS-CoV-2 as Compared with SARS-CoV-1. N Engl J Med. 2020;382(16):1564–7.

[15] Asadi S, Bouvier N, Wexler AS, Ristenpart WD. The coronavirus pandemic and aerosols: Does COVID-19 transmit via expiratory particles? Aerosol Sci Technol. 2020;0(0):1–4.

[16] Lin Y, Huang L, Nie S, Liu Z, Yu H, Yan W, et al. Knowledge, attitudes and practices (KAP) related to the pandemic (H1N1) 2009 among Chinese general population: a telephone survey. BMC Infect Dis. 2011;11:128.

[17] Zhong BL, Luo W, Li HM, Zhang QQ, Liu XG, Li WT, et al. Knowledge, attitudes, and practices towards COVID-19 among Chinese residents during the rapid rise period of the COVID-19 outbreak: a quick online cross-sectional survey. Int J Biol Sci. 2020;16(10):1745–52.

[18] ul Haq N, Hassali MA, Shafie AA, Saleem F, Farooqui M, Haseeb A, et al. A cross-sectional assessment of knowledge, attitude and practice among Hepatitis-B patients in Quetta, Pakistan. BMC Public Health. 2013;13:448.

[19] Yee I. The Philippines confirms its first case of coronavirus Atlanta: Cable News Network; 2020 [Available from: https://edition.cnn.com/asia/live-news/coronavirus-outbreak-01-30-20-intl-hnk/h_196b88233baee37963f367f8481eb226.

[20] BBC. Coronavirus: First death outside China reported in Philippines London: BBC; 2020 [Available from: https://www.bbc.com/news/world-asia-51345855.

[21] Tan L. Gov’t finalizes rules for areas under enhanced, general community quarantine Philippines: CNN Philippines; 2020 [Available from: https://cnnphilippines.com/news/2020/4/30/Philippines-ECQ-GCQ-lockdown-quarantine-guidelines-COVID-19.html.

[22] Gita-Carlos RA. Non-essential travel in ECQ, GCQ zones still banned: IATF-EID Philippines: Philippine News Agency; 2020 [Available from: https://www.pna.gov.ph/articles/1101520.

[23] Mubeen SM, Kamal S, Kamal S, Balkhi F. Knowledge and awareness regarding spread and prevention of COVID-19 among the young adults of Karachi. J Pak Med Assoc. 2020;70(Suppl 3)(5):S169–S74.

[24] Al-Mohrej A, Agha S. Are Saudi medical students aware of middle east respiratory syndrome coronavirus during an outbreak? J Infect Public Health. 2017;10(4):388–95.

[25] Galle F, Sabella EA, Da Molin G, De Giglio O, Caggiano G, Di Onofrio V, et al. Understanding Knowledge and Behaviors Related to CoViD-19 Epidemic in Italian Undergraduate Students: The EPICO Study. Int J Environ Res Public Health. 2020;17(10).

[26] Dalanon J, Ugalde RB, Catibod LD, Macaso JML, Okura K, Matsuka Y. Comparative analysis of education, awareness, and knowledge of dentists and physical therapists in the treatment of temporomandibular disorders. Cranio. 2020:1–8.

[27] Mohanna K. Teaching in the healthcare setting. Postgrad Med J. 2007;83(977):143–4.

[28] Wagner T. Incorporating Health Literacy Into English as a Second Language Classes. Health Lit Res Pract. 2019;3(3 Suppl):S37–S41.

[29] Hamza MS, Badary OA, Elmazar MM. Cross-Sectional Study on Awareness and Knowledge of COVID-19 Among Senior pharmacy Students. J Community Health. 2021;46(1):139–46.

[30] Prazeres F, Passos L, Simoes JA, Simoes P, Martins C, Teixeira A. COVID-19-Related Fear and Anxiety: Spiritual-Religious Coping in Healthcare Workers in Portugal. Int J Environ Res Public Health. 2020;18(1).

[31] Callueng C, Aruta J, Antazo BG, Briones-Diato A. Measurement and antecedents of national resilience in Filipino adults during coronavirus crisis. J Community Psychol. 2020;48(8):2608–24.

[32] Kimhi S, Eshel Y, Marciano H, Adini B. A Renewed Outbreak of the COVID-19 Pandemic: A Longitudinal Study of Distress, Resilience, and Subjective Well-Being. Int J Environ Res Public Health. 2020;17(21).

[33] Al-Hanawi MK, Angawi K, Alshareef N, Qattan AMN, Helmy HZ, Abudawood Y, et al. Knowledge, Attitude and Practice Toward COVID-19 Among the Public in the Kingdom of Saudi Arabia: A Cross-Sectional Study. Front Public Health. 2020;8:217.

[34] Mayol AV. Cebu City spends P2.5 million for ‘tuob’ kits Philippines: Inquirer; 2020 [Available from: https://newsinfo.inquirer.net/1305865/cebu-city-spends-p-2-5m-for-tuob-kits.

